# Transcranial magnetic stimulation induced pupil dilations can serve as a cortical excitability measure

**DOI:** 10.1101/2025.08.02.668303

**Authors:** Phivos Phylactou, Fraser MacRae, Athina Manoli, Nicolas Bouisset, Arnaud Duport, Afreen A. Awan, Xuhang Tong, Freek van Ede, David A. Seminowicz, Siobhan M. Schabrun

## Abstract

The application of non-invasive brain stimulation (NIBS) often relies on proxy estimates of cortical excitability (CE), such as the resting motor threshold (rMT), measured through transcranial magnetic stimulation (TMS). However, estimating the rMT is not always possible, as it requires an intact corticospinal pathway for neural signals to travel from the cortex to the periphery. To broaden the application of NIBS there is a need for additional CE measures. In three experiments combining TMS, transcranial direct current stimulation (tDCS), and eye-tracking, we measured TMS-induced pupil dilations as a potential CE proxy. We present Bayesian evidence (Experiment 1: BF > 9; Experiment 2: BF > 46; Experiment 3: BFs > 6) that TMS-induced pupil dilations serve as an objective CE proxy, with larger pupil size reflecting higher CE. We also show that these effects are not due to auditory, muscle, or sensory confounds. The introduction of this novel measure paves the way for a deeper understanding of CE by enabling objective NIBS measures beyond the motor cortex. Moving away from the reliance on the motor cortex would allow NIBS research and therapy to become more inclusive, allowing access to populations that are affected by damage along the corticospinal pathway.

---

Since its introduction 40 years ago [1], transcranial magnetic stimulation (TMS)—a safe, non-invasive brain stimulation (NIBS) technique—has been extensively used in both research and healthcare (for recent reviews see refs. [2, 3]). In contemporary practice, the application of TMS has predominantly relied on the primary motor cortex (M1), with motor evoked potentials (MEPs) serving as the primary objective proxy for cortical excitability (CE) (for examples see refs. [4, 5]).

However, the reliance of TMS on M1 creates two limitations. First, measuring MEPs requires an intact corticospinal pathway, thus restricting TMS applications to populations without damage along this pathway. Second, CE measures cannot be taken outside of the motor cortex objectively, since TMS MEPs can only be induced through M1 stimulation. Of note, though the phosphene threshold is an alternative CE measure [6], its estimation is subjective [7]. As such, there is a clear need for an additional objective proxy of CE.

Pupil size may serve as a potential CE proxy. Pupil dilation is driven by norepinephrine release that binds to receptors in the radial fibers of the dilator muscle. The main source of norepinephrine in the brain is the locus coeruleus (LC), a small nucleus in the brainstem that has one of the broadest projection patterns across the cortex [8].

Evidence from both animal [9] and human [10] studies has linked pupillary modulation to activity in the LC and its noradrenergic role (for a recent review see ref [11]). Moreover, with norepinephrine release acting on multiple cell targets (e.g., microglia, astrocytes, β2-adrenergic receptors) [8], the LC is thought to be pivotal for cortical plasticity, serving a functional role in neuromodulatory systems by dynamically facilitating or inhibiting neural networks [12]. Due to this noradrenergic mechanism it has been recently proposed that pupil size may serve as a marker of CE and cortical plasticity in general [11].

Early evidence from TMS studies suggests a link between pupil dilation and CE. For example, Niehaus et al. [13] showed that single trains of repetitive TMS (rTMS) with two, five, or ten pulses at 10 or 20 Hz induced pupil dilations, peaking around 1.5 seconds post rTMS, which were absent during sham rTMS. Further, in a small sample of four participants, they provided descriptive data suggesting that M1 TMS-induced pupil dilations may be dissociable across different TMS intensities. Along similar lines, Gimranov & Gimranova [14] illustrated that 1-minute of rTMS (2s of 10 Hz rTMS with 4s pause between trains; 200 pulses total) over occipital regions was followed by a decrease in pupil size, an effect that was absent during sham rTMS. Perhaps the strongest evidence supporting a link between pupil size and CE—as reflected through TMS—stems from a recent study by Weijs et al. [15], who measured MEPs in 15 participants who were trained to self-regulate their pupil size. Weijs et al. [15], while controlling for resting CE levels (no pupil self-regulation), showed that CE increased during upregulation (larger pupil size condition) compared to downregulation (smaller pupil size condition). Taken together, these findings demonstrate a link between CE and pupil size, suggesting that TMS may hold promise as a potential tool to induce and measure CE specific pupil responses.

Despite the promising nexus between CE and pupil size [11], this link currently stems from indirect evidence. For example, the primary aim of Niehaus et al. [13] was to investigate TMS effects on autonomous responses. Gimranov & Gimranova [14] investigated the role of occipital regions in adjusting pupil responses, while Weijs et al. [15] focused on measuring physiological responses to pupil self-regulation. Further, because of the different primary objectives of earlier TMS studies, alternative explanations of TMS-induced pupil responses have not been explored. As such, the overarching objective of this study was to directly investigate whether TMS-induced pupil dilations can serve as an objective proxy measure of CE, while controlling for possible confounders such as auditory, muscle, and sensory artefacts.

To achieve this objective, across three experiments, we combined NIBS with eye-tracking and measured TMS-induced pupil dilations in response to different stimulation intensities or before and after CE inhibition. Experiment 1 aimed to examine whether different TMS intensities result in distinct TMS-induced pupil dilations, while controlling for auditory artefacts with sham TMS. We predicted that dissociable pupil dilations would be evident across intensities and between active and sham TMS. The aim of Experiment 2 was to replicate the findings of Experiment 1 while controlling for additional confounders such as eye muscle artefacts through binocular eye-tracking and hand muscle artefacts through electromyography (EMG). In addition to replicating Experiment 1, we predicted that there would be no pupil lateralisation effects, indicating that TMS-induced pupil dilations are not due to spill-over stimulation of eye muscles. Additionally, we predicted that pupil responses to a given TMS intensity would not differ between low versus high hand muscle activity, demonstrating that TMS-induced pupil dilations are independent of peripheral muscle activity. Lastly, Experiment 3 employed transcranial direct current stimulation (tDCS) to investigate whether TMS-induced pupil dilations reflect CE inhibition. The aim of Experiment 3 was twofold: (i) to investigate if TMS-induced pupil dilations differ after CE inhibition compared to before, and (ii) to control for TMS sensory specific effects on pupil responses by maintaining TMS parameters consistent while shifting CE through a separate modality (i.e., tDCS). Our prediction was that TMS-induced pupil dilations would be smaller after inhibitory tDCS compared to before tDCS, suggesting that CE inhibition is reflected through TMS-induced pupil dilations and that pupil responses are not driven by sensory artefacts. The results from the three experiments supported our predictions.

## Results

Experiment 1 (*n* = 11) and Experiment 2 (*n* = 20) followed a similar within-participant, sham-controlled design (Fig. 1a). Specifically, 200 blink-free TMS pulses (100 active TMS trials and 100 sham TMS trials) were delivered with concurrent pupil recordings. Each condition (active vs sham; order counterbalanced) included five blocks of different intensities (80%-120% of rMT; 10% steps) delivered in random order. Experiment 3 (*n* = 28) followed a within-participant pre-post design, that included 80 TMS pulses at 120% of rMT in total, before, during, immediately after, and 10-minutes after a priming-test inhibitory tDCS protocol (Fig. 1b). In total, 59 healthy individuals participated across the three experiments.

**Fig. 1.**
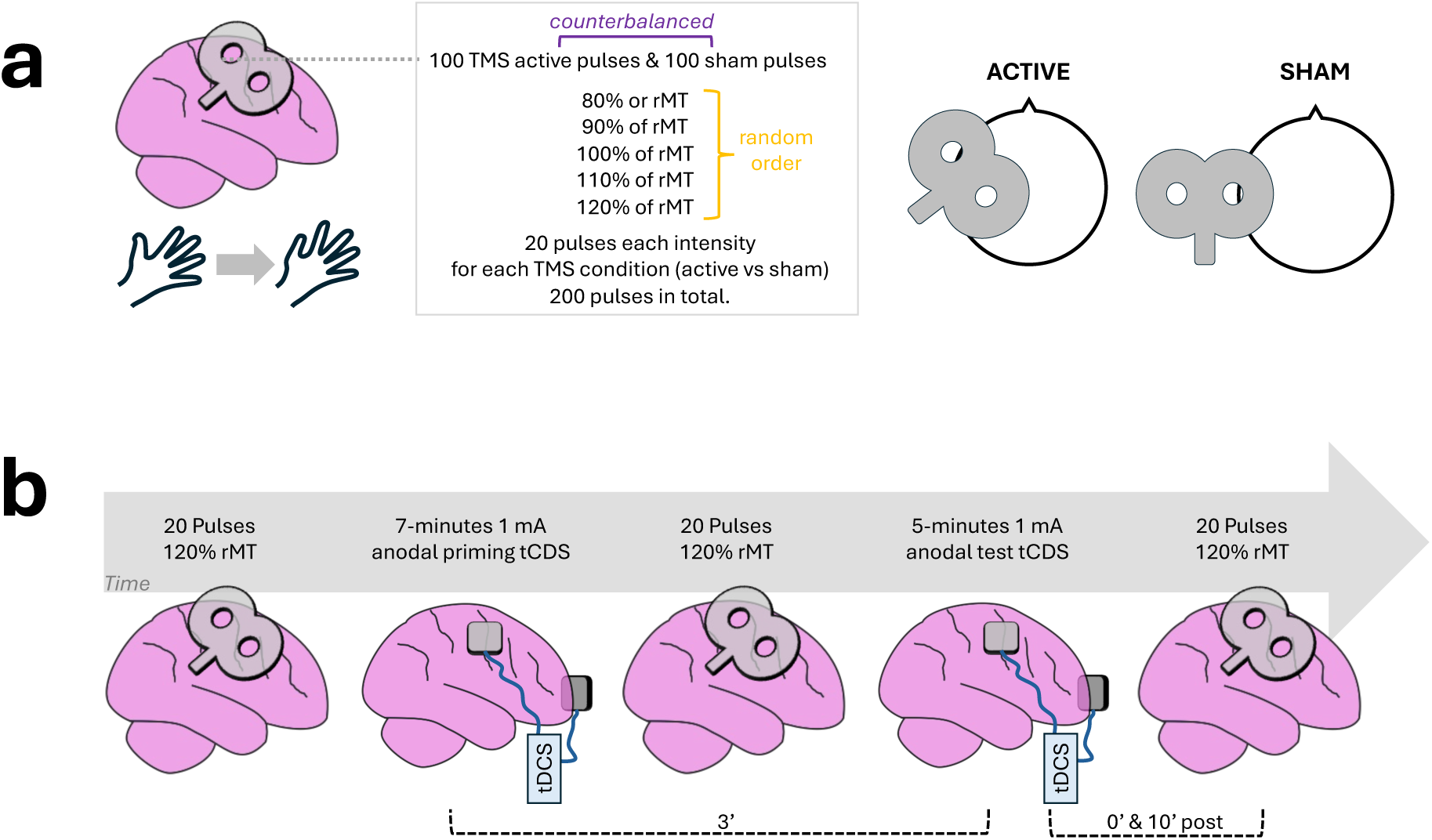
Overview of experimental designs for measuring TMS-induced pupil dilations. (**a**) Experiment 1 (*n* = 11) and Experiment 2 (*n* = 20) followed the same within-participant sham-controlled design. Participants completed 200 blink-free TMS trials. Five blocks of active TMS and five blocks of sham TMS were delivered in a counterbalanced order. Each block in each condition comprised a different TMS intensity, ranging from 80-120% (10% steps) of rMT in random order. To preserve the auditory artefact, during the sham TMS condition, the coil was placed with its side positioned on the hotspot, but the e-field targeted away from the cortex. (**b**) Experiment 3 (*n* = 28) had a within-participant pre-post design. Twenty blink-free TMS trials at 120% of rMT were collected before, during, immediately after, and 10-minutes after an inhibitory priming-test tDCS protocol. *Notes.* rMT: resting motor threshold; tDCS: transcranial direct current stimulation; TMS: transcranial magnetic stimulation.

### Experiment 1

In Experiment 1 (*n* = 11), pupil data recorded from the right eye revealed that active TMS resulted in consistent pupil dilations that peaked at approximately 1 second (*mean* = 1.06, *sd* = 0.31; Fig. 2a). A two (TMS condition: Active vs Sham) by five (Intensity: 80% vs 90% vs 100% vs 110% vs 120%) repeated measures Bayesian ANOVA (*R^2^ =* 0.67, 95% CI = [0.66, 0.68]) revealed that peak pupil dilations were dissociable for the TMS condition, Intensity, and the interaction of TMS condition and Intensity (*P*_(TMS+Intensity+TMS*Intensity|peak pupil)_ = 0.71, *BF*_model/model prior_ *=* 9.83; Fig. 2b). For reference, this winning model (TMS + Intensity + TMS*Intensity) superseded the null model by more than a ten-thousandfold (*BF*_model/null_ = 10387.14). Bayesian correlations echoed these results. In detail, a positive correlation was found between intensity and peak pupil in the active TMS condition (τ-β = 0.32, *BF =* 49.30), which was absent for sham TMS (τ-β = 0.13, *BF =* 0.46).

**Fig. 2.**
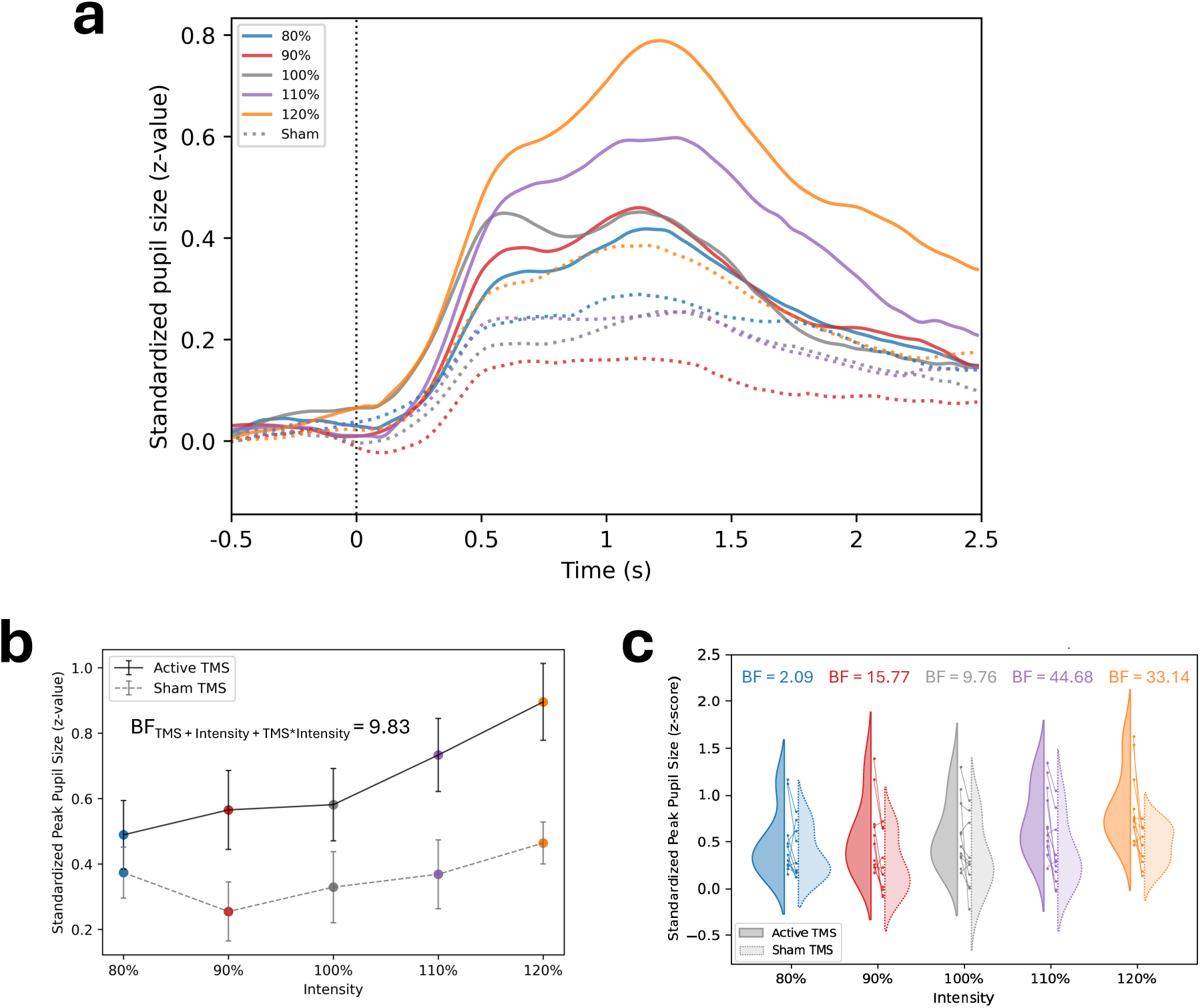
Results from Experiment 1. (**a**) Group averaged (n = 11) pupil traces relevant to the TMS pulse. (**b**) Active and sham TMS averaged peak pupil dilations per intensity condition. (**c**) Individual active versus sham TMS peak pupil dilations per intensity condition. *Notes.* Intensity is expressed as a percentage of the resting motor threshold. Pupil traces are smoothed with a Gaussian filter (σ = 1.5) for visualisation purposes only. Error bars depict standard error. TMS: transcranial magnetic stimulation.

Post-hoc pairwise comparisons, implemented through Bayesian Wilcoxon sign-ranked *t*-tests, resulted in Bayesian evidence supporting, with at least a two-fold evidence, that peak dilations were larger in the active compared to the sham TMS condition (Fig. 2c). Descriptive statistics and *t*-test results are provided in Supplementary Table S1(see also see also Supplementary Fig. S1). Further, evidence for within condition comparisons across intensities revealed that for the majority of contrasts in the active TMS condition, pupil dilations were dissociable between intensities, contrary to the sham TMS condition (Supplementary Table S2). Of note, in the sham TMS condition, differences across intensities were observed only when contrasting against the 120% condition.

Taken together, the proof-of-concept results from Experiment 1 show that TMS-induced pupil dilations differ between intensities, with higher TMS intensities resulting in larger pupil dilations. These effects are absent during sham TMS, suggesting that TMS-induced pupil dilations are not driven by auditory artefacts. To confirm these results and to control for possible alternative explanations, we replicated the design in a second experiment with a larger sample size, while taking additional measures to facilitate control analyses.

### Experiment 2

In Experiment 2, a larger sample (*n* = 20) underwent similar procedures as in Experiment 1. To enable the exploration of alternative explanations of the results from Experiment 1, Experiment 2 included electromyography (EMG) recordings as well as binocular eye-tracking. Further, a random jitter (see *Materials and Methods*) was introduced between trial initiation and the TMS pulse to control for anticipation effects. Additionally, TMS procedures in Experiment 2 utilized stereotactic neuronavigation.

*Replication.* Results replicated the findings of Experiment 1. Binocular pupil data (averaged across the left and right pupil) showed a consistent TMS-induced pupil dilation peaking around 1 second (*mean* = 1.08, *sd* = 0.31; Fig. 3a). As earlier, a two (TMS condition: Active vs Sham) by five (Intensity: 80% vs 90% vs 100% vs 110% vs 120%) repeated measures ANOVA (*R^2^ =* 0.61, 95% CI = [0.61, 0.61]) was implemented. Evidence supported that peak pupil dilations were different across TMS condition, Intensity, as well as the interaction of TMS condition and Intensity (*P*_(TMS+Intensity+TMS*Intensity|peak pupil)_ = 0.92, *BF*_model/model prior_ *=* 46.99; Fig. 3b), with this model (TMS + Intensity + TMS*Intensity) superseding the null model (*BF*_model/null_ = 3696.12). These effects were reflected through correlation analyses, that provided evidence for a correlation between intensity and peak pupil dilation for the active (τ-β = 0.21, *BF =* 14.55) but not the sham condition (τ-β = 0.01, *BF =* 0.13).

**Fig. 3.**
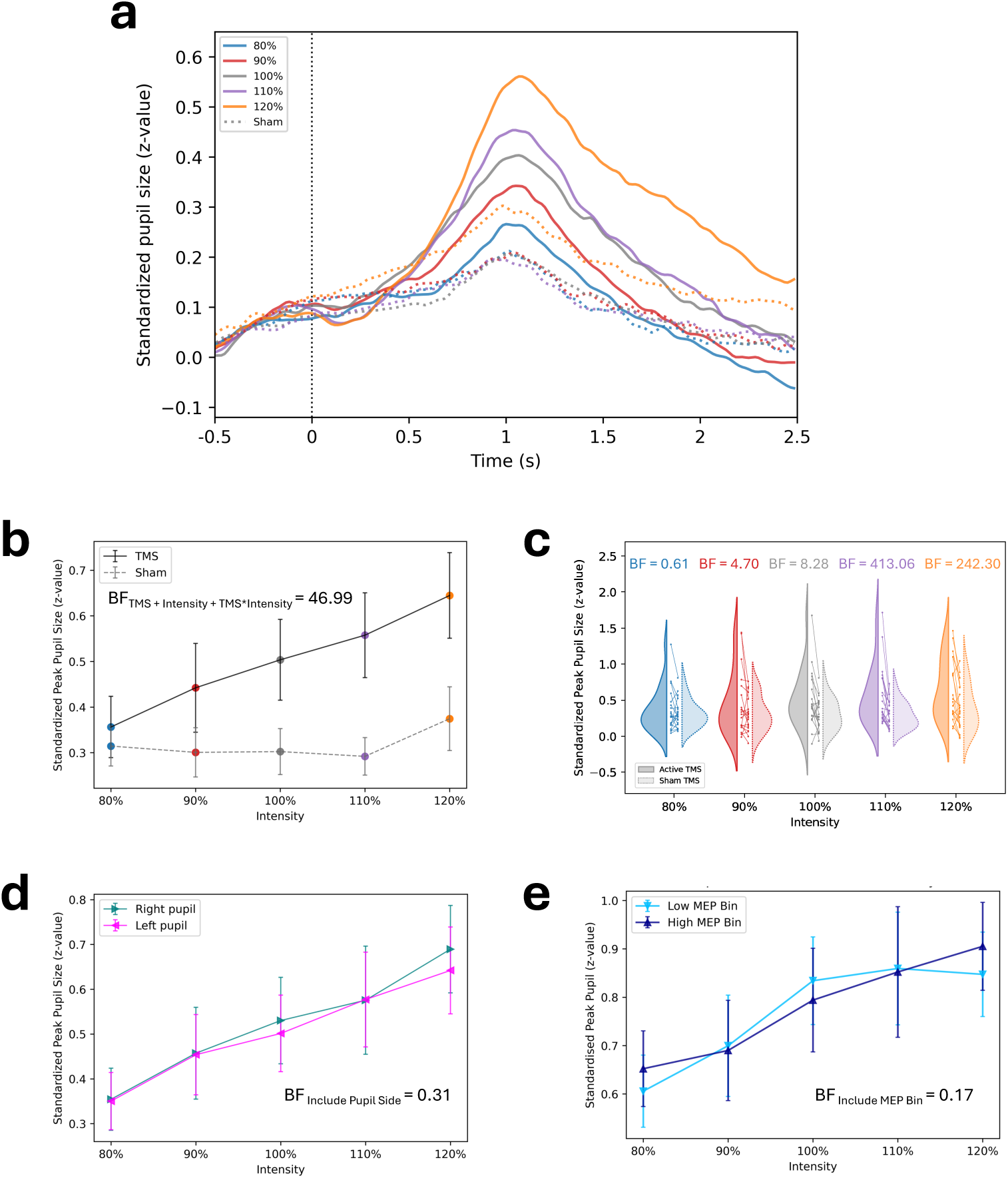
Results from primary analyses in Experiment 2. (**a**) Group averaged (n = 20) pupil traces relevant to the TMS pulse. (**b**) Active and sham TMS averaged peak pupil dilations per intensity condition. (**c**) Individual active versus sham TMS peak pupil dilations per intensity condition. (**d**) Comparisons between left versus right pupil in the active TMS condition. (**e**) Pupil size comparisons following within-participant median split grouping of each condition’s MEP data (Low MEP vs High MEP bin). *Notes.* Intensity is expressed as a percentage of the resting motor threshold. Pupil traces are smoothed with a Gaussian filter (σ = 1.5) for visualisation purposes only. Error bars depict standard error. MEP: motor evoked potential; TMS: transcranial magnetic stimulation.

Pairwise analyses revealed that in all but the 80% condition, active TMS resulted in larger pupil responses compared to sham TMS (Fig. 3c; Supplementary Table S3; see also see also Supplementary Fig. S1). Within-condition intensity comparisons provided supporting evidence that pupil peaks were different across intensities in the active TMS condition, but not in the sham TMS condition (Supplementary Table S4). Similar to Experiment 1, differences across intensities in the sham TMS condition were noticed only in comparison with the 120% condition.

*Muscle confounds.* An alternative explanation for the TMS-induced pupil dilations can be attributed to muscle confounds. For example, pupil dilations may not be due to cortical TMS per se, but instead, due to spill-over stimulation of the pupil’s dilator muscle. In such a case, lateralisation effects should be evident, with the left pupil, which is closer to the TMS coil, producing larger peaks post TMS compared to the right eye. We tested this prediction using a two (Pupil Side: Left vs Right) by five (Intensity: 80% vs 90% vs 100% vs 110% vs 120%) repeated measures ANOVA for the active TMS condition (*R^2^ =* 0.66, 95% CI = [0.66, 0.67]; Fig. 3d). The ANOVA showed that the model with the highest probability given the data, was the model including only the Intensity factor (*P*_(Intensity|peak pupil)_ = 0.67, *BF*_model/model prior_ *=* 8.23). Adding the Pupil Side factor to this model reduced the probability of the model (*P*_(Pupil Side+Intensity|peak pupil)_ = 0.27), with the Intensity model superseding the Intensity + Pupil Side model (*BF*_Intensity/Intensity+Pupil Side_ = 2.49). Further, the exclusion probability of the Pupil Side factor given the data was *P_(exclude Pupil Side|peak pupil)_* = 0.68 (*BF_include_* = 0.31). These findings were further supported by evidence in favor of the null in paired *t*-tests that tested for lateralisation differences for each condition separately (Supplementary Table S5; Supplementary Fig. S2).

In addition to eye-muscle confounds, another alternative explanation may lie in FDI muscle artefacts. For instance, considering the increased FDI activity given higher intensities under active TMS, it is possible that TMS-induced pupil dilations reflect peripheral muscle movement rather than cortical stimulation. To explore this alternative, we separated the peak pupil data in each intensity condition into two data bins, based on the corresponding EMG activity in each trial. The separation was based on a median split analysis performed on the MEP data, separately for each intensity in the active TMS condition, resulting in a Low MEP bin and a High MEP bin for each intensity and condition. A repeated measures ANOVA was performed, testing a two (MEP Bin: Low vs High) by five (Intensity: 80% vs 90% vs 100% vs 110% vs 120%) model (*R^2^ =* 0.71, 95% CI = [0.65, 0.76]; Fig. 3e). The model including only the Intensity factor resulted in the highest probability given the data (*P*_(Intensity|peak pupil)_ = 0.80, *BF*_model/model prior_ *=* 15.89). Model probability was reduced when accounting for the MEP bin (*P*_(MEP Bin+Intensity|peak pupil)_ = 0.18), with the Intensity model surpassing the Intensity + MEP bin one (*BF*_Intensity/Intensity+MEP Bin_ = 4.44). The MEP Bin factor exclusion probability was *P_(exclude MEP Bin|peak pupil)_* = 0.80 (*BF_include_* = 0.17). With the exception of the 120% condition for which there was inconclusive evidence, paired *t*-test resulted in evidence favoring the null, when comparing the Low MEP versus High MEP bins in each condition (Supplementary Table S6; Supplementary Fig. S2).

*Relationship of MEPs and Pupil data.* As anticipated, MEP data were also dissociable between active and sham TMS, and across intensities in the active TMS condition, as supported by evidence from a repeated measures ANOVA (*P*_(TMS+Intensity+TMS*Intensity|MEP)_ = 1, *BF*_model/model prior_ *=* 2.62×10^12^). Further, MEPs in the active TMS condition correlated with TMS intensity (τ-β = 0.53, *BF =* 2.48 x10^12^). Even though both MEPs and TMS-induced pupil dilations were independently correlated with TMS intensity (Fig. 4a), MEPs and TMS-induced pupil dilations were not correlated with each other (τ-β = 0.01, *BF =* 0.13; Fig. 4b).

**Fig. 4.**
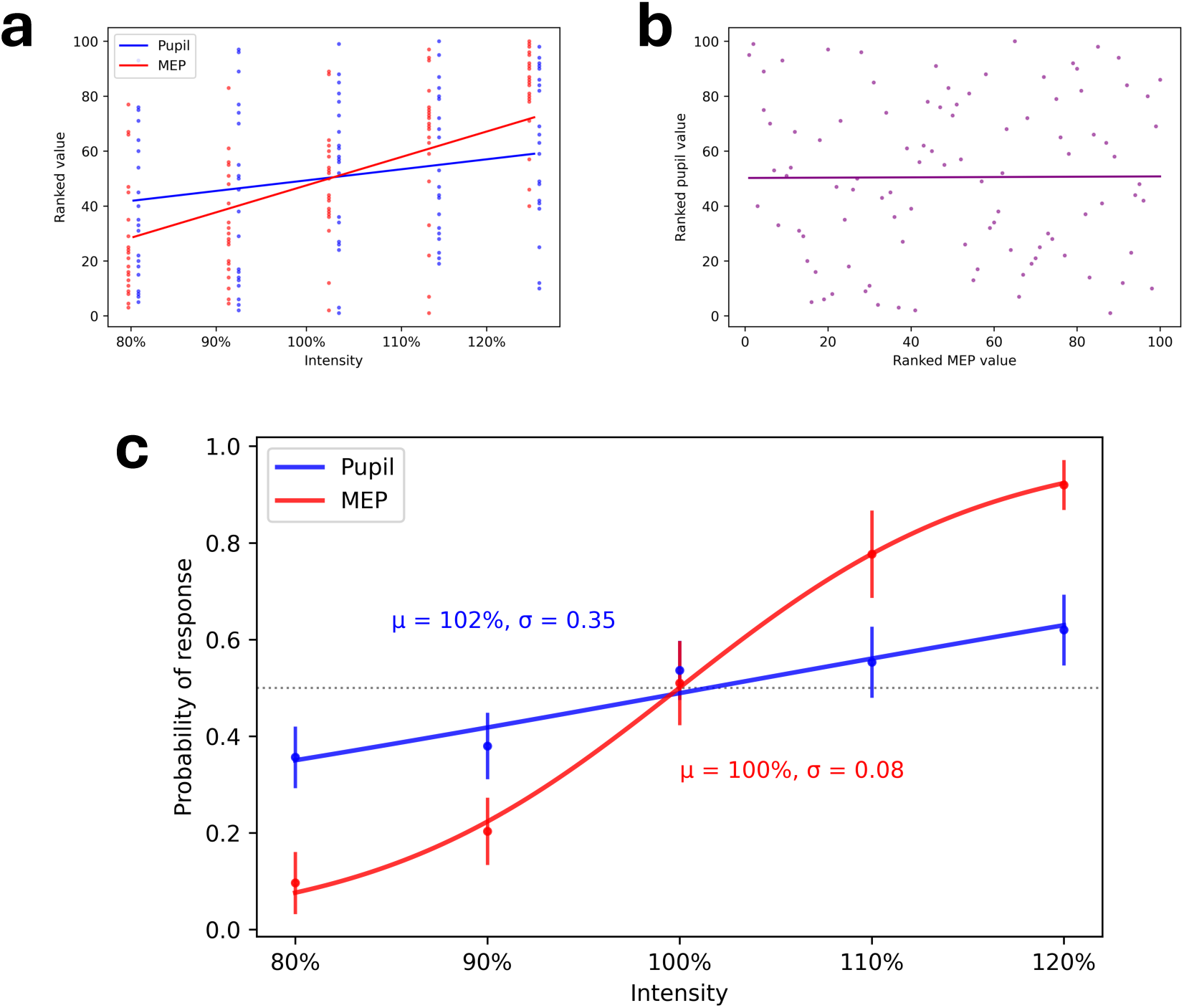
Exploratory analyses investigating the relationship between TMS induced MEPs and TMS-induced pupil dilations. (a) Correlations of MEPs (red) and peak pupil dilations (blue) with TMS intensity. (b) Absence of correlation between MEPs and peak pupil dilations. (c) Psychophysical logistic curve of MEP (red) and peak pupil (blue) data as a function of TMS intensity. *Notes.* Intensity is expressed as a percentage of the resting motor threshold. Error bars depict standard error. MEP: motor evoked potential; TMS: transcranial magnetic stimulation.

The lack of correlation between MEPs and TMS-induced pupil dilations, could be attributed to the different threshold sensitivity of each measure. As an exploratory approach to investigate this possibility, psychophysical logistic curves were fitted to both the MEP and pupil data. We used a probabilistic curve to facilitate comparisons across the two measures. In detail, the data were fitted using the equation *θ =* 1 / (1+ *e*^-(χ-μ)σ^), where χ is the probability of response calculated for each condition, μ is the curve shift (point of inflection), and σ is the curve slope. For MEPs, the threshold to calculate the probability of response was set to 50 μV, following our rMT procedure (see *Materials and Methods*). For pupil data, the probability of response threshold was set as the median pupil peak value (*z* = 0.65), calculated across all participants for all trials in the active TMS condition. The curves were successfully fitted to 15 participants; curve fitting failed for the MEP data of two participants (μ_MEP_ < 0) and for the pupil data of three participants (μ_pupil_ < 0). The resulting functions (Fig. 4c) provide some insight as to the potential sensitivity differences between MEPs and pupils. Specifically, the group-level curves show similar shift μ for pupils (μ_pupil_ = 102%) and MEPs (μ_MEP_ = 100%), with the curve slope σ being shallower for pupils (σ_pupil_ = 0.35) compared to MEPs (σ_MEP_ = 0.08). As an additional exploratory analysis, we compared individual-level (*n* = 15) curve shift and slope between pupil and MEP functions, using paired *t-*tests with a Cauchy distribution centered on 0 and with a scale *r* = 0.707. For curve shift μ, evidence suggested that there were no differences (*BF_10_* = 0.29) between pupil curve shift (mean μ_pupil_ = 111.22, *sd* = 45.58) and MEP curve shift (mean μ_MEP_ = 97.98, *sd* = 22.01). For curve slope σ, evidence (*BF_10_* = 364.01) suggested steeper curves for MEP (mean σ_MEP_ = 0.05, *sd* = 0.05) compared to pupil (mean σ_pupil_ = 0.37, *sd* = 0.28). These results provide some exploratory evidence suggesting that MEPs and TMS-induced pupil dilations likely have different sensitivity profiles.

Overall, the results in Experiment 2 replicated the findings of Experiment 1 with additional evidence supporting the absence of pupil lateralization effects, as well as the absence of pupil peak differences based on FDI activity. Correlational and psychophysical analyses suggested that pupil peaks and MEPs have different sensitivities as proxy measures of CE.

### Experiment 3

The aim of Experiment 3 (*n =* 28) was twofold: (i) to investigate if TMS-induced pupil dilations differ after compared to before CE inhibition, and (ii) to further explore whether TMS effects are due to sensory confounds (e.g., pupil dilations driven by different sensations on the skull due to differences in TMS intensity). To achieve these aims, we introduced a priming-test tDCS protocol, to decrease CE [16]. If TMS-induced pupil dilations reflect CE, then pupil peaks should be smaller following inhibition as compared to baseline pupil peaks. In parallel, since TMS parameters are consistent and a different modality is utilised to inhibit CE, if TMS-induced pupil dilations are due to sensory confounds (i.e., the findings in Experiment 1 and 2 are because of increased sensory artefacts due to higher TMS intensities), then pupil peaks will remain unchanged before versus after tDCS.

To confirm the induction of tDCS inhibition, we compared MEP peak-to-peak amplitudes before and after tDCS (Fig. 5a). Compared to baseline, as expected (see *Materials and Methods*), MEP peak-to-peak amplitude was decreased 10 minutes post tDCS (*BF_10_* = 4.50), whereas evidence favored the null for differences in MEPs immediately post tDCS (*BF_10_* = 0.71; Fig. 5b). As for the pupils, as in Experiment 1 and Experiment 2, consistent TMS-induced pupil dilations were evident, peaking at 1 second (*mean* = 1.04, *sd* = 0.32; Fig. 5c). Compared to baseline, TMS-induced pupil dilations were smaller both immediately after (*BF_10_* = 57.60) and 10-minutes after tDCS (*BF_10_* = 6.03). Descriptive statistics for both MEP and pupil data are presented in Supplementary Table S7.

**Fig. 5.**
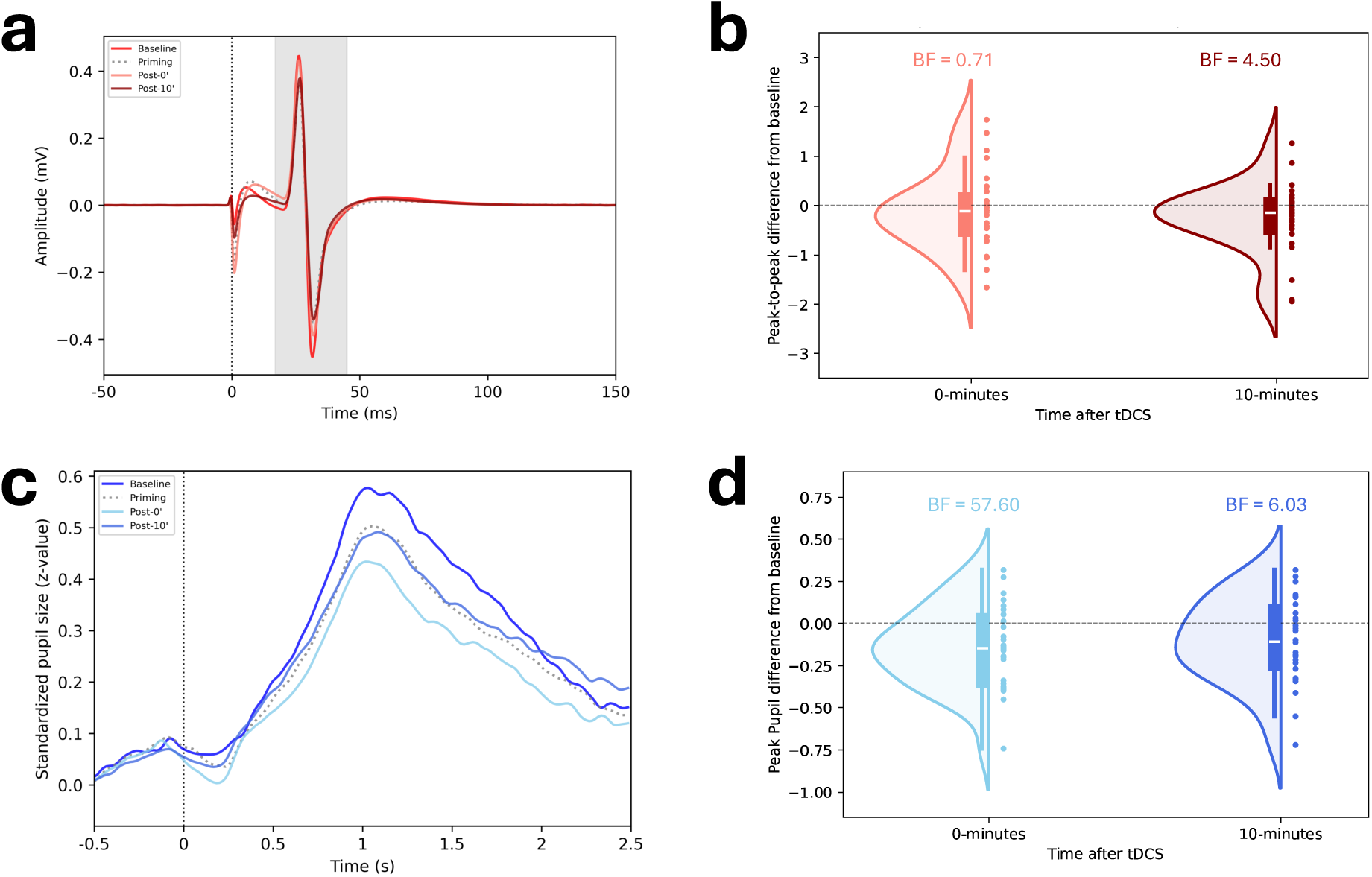
Results of Experiment 3. (a) MEP traces before, during, immediately after, and 10 minutes after the induction of a priming-test tDCS inhibitory protocol. (b) Difference plots comparing baseline MEP peak-to-peak amplitudes with those immediately after and 10 minutes after tDCS. (c) Group averaged (n = 28) pupil traces relevant to the TMS pulse, before, during, immediately after, and 10 minutes after the induction of a priming-test tDCS inhibitory protocol. (d) Difference plots comparing baseline peak TMS-induced pupil dilation with those immediately after and 10 minutes after tDCS. *Notes.* MEP: motor evoked potential; tDCS: transcranial direct current stimulation. TMS: transcranial magnetic stimulation.

Complementary to the findings from Experiments 1 C 2, Experiment 3 indicates that the pupil reflects the reduction of CE through decreased TMS-induced pupil dilations. In addition, changes to the pupil response following tDCS inhibition support the notion that TMS-induced pupil dilations are not due to TMS sensory artefacts.

## Discussion

Across three experiments, this study presents evidence suggesting that TMS-induced pupil dilations can serve as a proxy CE measure. In all three experiments, consistent TMS-induced pupil dilations were observed, which peaked at 1 second post-TMS. Experiment 1 illustrated that these TMS-induced pupil dilations were dissociable across TMS intensities, with stronger TMS resulting in larger pupil dilation peaks. Further, TMS-induced pupil dilations were distinct from sham TMS, in support of the notion that pupil responses were not driven by auditory artefacts. Experiment 2 replicated these findings and showed that TMS-induced pupil dilations were not due to eye-muscle stimulation, supported by evidence against pupil lateralisation effects. Additionally, results supporting no difference in pupil dilation peaks between low versus high MEP trials demonstrated that TMS-induced pupil dilations were not due to peripheral muscle activity. When it comes to the relationship between TMS-induced pupil dilations and MEPs, exploratory findings suggest the two measures likely have different sensitivity thresholds.

Lastly, Experiment 3 found that, following inhibitory tDCS, TMS-induced pupil dilations were smaller immediately after and 10 minutes after tDCS—as compared to before tDCS— thus reflecting CE inhibition. In parallel, since TMS parameters were unchanged in Experiment 3, these data also suggest that TMS-induced pupil dilations were not due to sensory specific TMS effects.

Contrary to previous work showing that pupil dilations in response to TMS peaked at 1.5 seconds [13], our results showed a consistent peak at 1 second. This discrepancy may be due to the use of different stimulation parameters. In detail, contrary to the single pulse TMS that was used here, Niehaus et al. [13] used rTMS trains and it is unclear how latency was calculated in relation to the various rTMS train conditions. Another explanation may lie in potential differences in luminance conditions. Though it is not known what the lighting conditions were in Niehaus et al.’s study [13], we showed consistent peak latencies in two different isoluminant conditions, at 400 lux (Experiment 1) and 300 lux (Experiment 2 C Experiment 3). Alternatively, the developments in eye-tracking technology in the past decades (see ref. [11]), could also explain the latency differences compared with earlier studies.

The dissociable pupil responses across intensities provide evidence for the use of TMS-induced pupil dilations as a CE proxy. In a small sample (*n* = 4), Niehaus et al. [13] previously provided descriptive data suggesting that pupil dilations may differ between TMS intensities. Our findings, from two independent experiments, provide Bayesian evidence that stronger TMS pulses result in larger pupil dilations that are distinct across intensities. Because of these dissociable pupil responses, it is possible that TMS-induced pupil dilations could function as an additional objective proxy measure of CE, serving a similar role as MEPs.

When exploring the relationship between TMS-induced pupil dilations and MEPs, we found evidence of absence for a correlation, even though both measures correlated with TMS intensity independently. This could be attributed to each measure having a different sensitivity threshold for CE. Indeed, our psychophysical curve functions suggest that TMS-induced pupil dilations have a lower sensitivity threshold (shallower slope) compared to MEPs (steeper slope). Although, it should be noted that the current study was not tailored to fit such functions, hence these analyses should be considered exploratory in nature. Such threshold sensitivity differences are aligned with previous comparisons between TMS measures, showing that the relationship across thresholds is complex. Recent meta-analytic work [6] demonstrated that even though the rMT and the phosphene threshold are correlated, when controlling for different TMS parameters, the linear relationship between the measures changes. This meta-analysis also showed that even though the thresholds were correlated, they were quantitatively different [6]. This illustrates that relying on M1 for TMS application, as the only available objective CE measure, may not necessarily provide a representative CE metric, considering also that it is by nature arbitrary (see ref. [6]). Introducing a second measurable proxy, such as the pupil size, would facilitate objective comparisons (contrary to phosphenes which are subjective [7]) and a deeper understanding of CE.

To provide further evidence that TMS-induced pupil dilations reflect CE and are not due to TMS sensory artefacts, we inhibited CE using tDCS. Using an inhibitory priming-test tDCS protocol [16–18], we showed that under consistent TMS parameters, TMS-induced pupil dilations were smaller following tDCS inhibition, compared to before tDCS. Inhibition was confirmed through a reduction in the MEP size 10 minutes post tDCS, which is the expected inhibition peak as reflected through MEPs [16, 18] (for a review see ref. [19]). A reduction of TMS-induced pupil dilations was evident both immediately after and 10 minutes after tDCS. Earlier work that applied rTMS over visual regions to inhibit CE, concluded that pupil size reductions following neuromodulation were most evident immediately after stimulation [14]. Considering the possibility of different sensitivity thresholds, together with the fact that MEPs are also contaminated by other muscle, spinal, and peripheral signals, it is likely that CE inhibition is reflected earlier through TMS-induced pupil dilations compared to MEPs.

A potential mechanism underpinning TMS-induced pupil dilations is noradrenergic LC activity. Though some researchers consider pupil responses as a direct measure of LC driven noradrenergic activity (for a recent review see ref. [20]), our findings, as well as those of earlier studies [13–15], cannot be directly attributed to LC activity. Along similar lines, evidence linking CE, as measured through TMS MEPs, with LC activity is also indirect. For example, increased CE in healthy participants was shown following pharmacological interventions of noradrenaline reuptake inhibitors that are used to increase norepinephrine in the brain [21–23]. Similarly, decreased CE in healthy participants was found after interventions with anti-noradrenergic drugs (i.e., α2-adrenergic agonist guanfacine) [24]. To study the neural architecture of TMS-induced pupil dilations, subsequent work should combine TMS and eye-tracking with concurrent neuroimaging (e.g., functional magnetic resonance imaging).

Our findings should be considered alongside the study’s limitations. First, even though TMS-induced pupil dilations seem to reflect CE with a consistent pupil trace profile across experiments, it remains unknown whether this response is stable across time within individuals. Future work should examine the reliability of the measure across the short-(e.g., minutes/hours) and long-(e.g., days/weeks) term. Second, even though we showed that pupil responses were not confounded by FDI muscle artefacts, it is possible that the TMS effects are specific to physiological responses (see ref. [25]). As such, TMS-induced pupil responses outside of M1 should be explored. Lastly, even though Experiment 3 used a tDCS protocol that has been shown to be reliable [16, 19] and inhibition was confirmed through the expected MEP aftereffects, it lacked a sham control condition. Replication with sham controlled protocols as well as with different montages (i.e., excitatory NIBS) will strengthen the interpretation of our findings.

Overall, we demonstrate that TMS-induced pupil dilations can serve as a CE proxy. In three experiments, we show that pupil responses following TMS are distinct across TMS intensities and before versus after CE inhibition. Introducing a new measurable proxy of CE enables comparisons with M1 derived TMS measures and paves the way for an in-depth understanding of CE and cortical plasticity. Finally, it offers the potential for making TMS more inclusive, by enabling TMS applications for individuals with damage anywhere along the corticospinal tract (e.g., MEP negative [26]), and more effectual by facilitating objective TMS measures outside of M1.

## Materials and Methods

In its whole, the study included 59 participants (28 females, 31 males; 27 women, 31 men, 1 non-binary; age mean = 25.73, *sd* = 5.83, min-max = [18, 49]), who provided written informed consent for their participation. All participants self-reported as right-handed and had normal or corrected-to-normal vision. To be eligible for the study, participants were screened for TMS safety based on recommended guidelines by Rossi et al. [27, 28]. The questions used for eligibility screening are provided in Supplementary Table S8, and included screening for physical, neurological, and mental health. Eligible participants were not asked to refrain from their regular routines (e.g., abstain caffeine, nicotine, or motor activity). The study received ethical approval from Western University’s Health Sciences Research Ethics Board (WREM ID: 126067) and the LAWSON Research Institute at St. Joseph’s HealthCare Ethics Board (ReDA ID: 15249).

Details about the methodological procedures are presented separately for each experiment in the following subsections. The reporting of the described procedures adheres to the TMS Reporting Assessment Tool (TMS-RAT v0.3, https://tms-rat.org). No part of the current study was pre-registered.

### Experiment 1

Experiment 1 served as a proof-of-concept study. Its purpose was to investigate whether TMS-induced pupil responses are distinct across intensities and between active and sham TMS.

#### Participants

Eleven healthy participants (8 females, 3 males; age mean = 26.73, *sd* = 3.35, min-max = [20, 32]) took part in Experiment 1. Considering the proof-of-concept purpose, the experiment comprised a convenience sample without an a priori sampling plan.

#### Apparatus

A biphasic MagStim Rapid^2^ with a Plus^1^ module (MagStim, Whitland, Wales, UK) was used for TMS. A 70 mm outer diameter figure-of-eight MagStim D70 alpha coil (P/N: 4150-00) was connected to the TMS device to deliver the pulses. All experimental procedures were designed and controlled using PsychoPy v2024.2.1 [29]. The TMS device and pulse delivery were controlled with the MagPy TMS v1.4 package [30]. Pupil data were recorded from the right eye, using an EyeLink 1000 Plus (SR Research, Ottawa, Canada) desktop eye-tracker, recording at 60 Hz through PsychoPy using the EyeLink API. The eye-tracker was set for a 21.5″ monitor with a 60 Hz refresh rate. During eye-tracking data collection, a chin and forehead rest was used to maintain participants’ heads stable, at a 60 cm viewing distance from the screen. All procedures took place in a single 2-hour session under isoluminant conditions (∼400 lux).

#### Hotspot determination

Throughout Experiment 1, TMS was delivered over right M1, at the optimal position to target the left FDI muscle. Participants were seated comfortably on a chair, with both hands resting on a pillow, and were asked to maintain a relaxed state. For all stimulations, the coil was placed at a 45° angle from the sagittal plane with the coil handle pointing posteriorly in relation to the head. This coil placement is meant to deliver a current with a posterior-anterior direction during the first phase and an anterior-posterior direction during the second phase of the biphasic pulse. To determine the FDI hotspot, participants were fitted with a tight cap marked with the 10-20 electroencephalography system scalp locations. The coil was moved around a 5 cm by 5 cm grid centred on the C4 scalp location, and three TMS pulses separated by at least five seconds were delivered at each position at an intensity set at 60% of maximum stimulator output (MSO) until FDI muscle twitches were visible. If no muscle twitches were elicited, the intensity was increased by 5% and the procedure was repeated until FDI twitches were induced. When the optimal FDI position was localised, stickers were used to trace the optimal coil positioning on the cap. After marking the cap, the coil was placed back on the marked position, and three TMS pulses were delivered to confirm the hotspot.

#### Resting motor threshold

The rMT was estimated as the MSO intensity required to induce visible twitches of the FDI muscles in approximately five out of 10 stimulations. To calculate this, a flexible staircase procedure was employed following, in order, steps of 4, 2, and 1 in absolute percentage units. The initial MSO intensity was set 5% below the intensity used to identify the hotspot. A visible muscle twitch would result in a decrease in intensity (based on the step), while a lack of a muscle twitch would result in an increase in intensity. Following three consecutive intensity reversals, up to 10 pulses (at least five seconds apart) were delivered at the given intensity. If FDI twitches were visible with a success rate of 100% or 0% after 4 pulses, the intensity was increased or decreased by 2±1 units, respectively, and TMS pulses were repeated until the threshold criterion (∼5 out of 10) was reached.

#### TMS-induced pupil size recordings

To record TMS-induced pupil dilations, participants were instructed to fixate on the centre of a black screen. Each trial had a duration of five seconds, during which participants were instructed to keep their eyes open and refrain from blinking. Trials were monitored online, and trials contaminated with eye-blinks were marked for exclusion and were replaced by new trials. Each trial was participant initiated to enable rest and blinking, as well as to maintain participant focus and attention. To initiate a trial, participants pressed the spacebar on a keyboard with their right hand. Two seconds after trial initiation a TMS pulse was delivered, followed by a three second delay. Pupil size was recorded for the whole 5 second duration of each trial.

#### Design and procedure

Experiment 1 followed a counterbalanced sham-controlled design. Following hotspot and rMT determination, the coil was placed over the hotspot and was secured in position using a mechanical arm in combination with the chin and forehead rest. Next, participants underwent five blocks of either sham or active TMS, followed by five blocks of the opposite condition, with the order counterbalanced across participants. For the sham TMS condition, the coil was placed with its side touching the hotspot and the focal point of the magnetic e-field facing towards the floor at the side of the head. This approach was preferred over a sham coil to preserve the TMS auditory artefact.

Participants were blinded to the sham design of the study but were told that TMS would be delivered at two different positions. Before the start of each condition (active and sham) a 9-point eye-tracker calibration was performed. In each block, TMS was delivered at a different intensity at either 80%, 90%, 100%, 110%, or 120% of the rMT. For each participant and for each condition, the order of the blocks was delivered randomly. For each block, participants had to complete 20 blink free trials, resulting in 200 blink free trials in total (100 active TMS trials, 100 sham TMS trials). Trials with TMS induced blinks (i.e., blinks with or immediately after the TMS pulse) were not rejected. The design of Experiment 1 is illustrated in Fig. 2a.

### Experiment 2

Experiment 2 was designed to replicate the findings of Experiment 1 in a new, larger participant cohort. Further, additional measures were included (EMG and binocular eye-tracking) to enable the investigation of potential alternative explanations of the effects found in Experiment 1.

#### Participants

Twenty healthy individuals (11 females, 9 males; age mean = 24.65, *sd* = 5.39, min-max = [18, 39]) participated in Experiment 2. The sample size for Experiment 2 was determined following a Bayesian Sequential Analysis approach [31], similar to previous TMS research [32]. Specifically, the sampling plan that we followed comprised an initial round of our primary analysis (see *Data Analysis* subsection) after collecting data from 20 participants. If our decision threshold (*BF* > 3) was not reached, four more participants were to be added before the next analysis round, and this procedure was to be repeated, until the decision threshold was reached. Here, the decision threshold was reached at the first round of analysis (*n* = 20).

#### Apparatus

As in Experiment 1, a biphasic MagStim Rapid^2^ Plus^1^ TMS system was used. TMS was delivered using a 70 mm outer diameter figure-of-eight MagStim D70 remote controlled coil (P/N: 3190-00). The experiment was controlled using PsychoPy v2024.2.4 [29] and the MagPy TMS v1.4 package [30]. The eye-tracker was configured for a 24″ monitor set at a 60 Hz refresh rate. Eye-tracking was recorded binocularly, using a Tobii Pro Spark (Tobii, Stockholm, Sweden) with a 60 Hz sampling rate, collected through PsychoPy using the Tobii API. The eye-tracker and screen were set at a 60 cm viewing distance. For Experiment 2, a Brainsight stereotactic neuronavigation system (Rogue Research, Montreal, Canada) was used to guide TMS coil placement and maintain head stability. Additionally, Brainsight’s Model 3 built-in MEP-box amplifier (Rogue Research, Montreal, Canada) was used to record EMG activity from the FDI. All procedures took place in a single 2-hour session in an isoluminant environment (∼300 lux).

#### Electromyography

EMG was recorded at 3000 Hz, with a bandwidth of 16-550 Hz and a 60 Hz notch filter, from the FDI muscle of the right hand using 30 mm by 24 mm pre-gelled Ag-AgCl surface electrodes. The electrodes were placed in a belly-tendon montage, with the negative electrode centred on the muscle belly and the positive electrode placed two cm laterally (towards the direction of the metacarpophalangeal joint). The ground electrode was placed over the ulnar styloid process. Before electrode placement, the skin area under each electrode position was cleaned using gauze soaked with 70% isopropyl rubbing alcohol, which after air drying, was followed by application of electrode skin preparation gel. During TMS, trials with background EMG activity (i.e., FDI muscle not at rest), which were monitored online, were replaced by new trials.

#### Hotspot determination

Participant set-up and coil orientation were identical to Experiment 1. Contrary to Experiment 1, TMS was delivered on the left M1, targeting the right FDI muscle. To determine the FDI hotspot in Experiment 2, participants’ heads were co-registered on a 1 by 1 by 1 mm^3^ averaged T1-weighted anatomical magnetic resonance image (MNI152 space). This co-registration was used throughout the experiment. To localise the hotspot, the coil was moved within a 5 cm by 5 cm grid centered on MNI coordinates x = -42, y = -19, z = 59. The hotspot was determined as the coil position resulting in the largest FDI MEPs. As in Experiment 1, hotspot localisation was conducted starting at a 60% MSO intensity, with increments of 5% in cases of no visible MEPs within the grid. The hotspot location was then marked in the neuronavigation software and was used for all subsequent TMS delivery, with an error tolerance of 2 mm for target distance and 3° for coil orientation and tilt.

#### Resting motor threshold

For Experiment 2, an uninformed psychophysical adaptive Maximum-Likelihood algorithm was employed to estimate rMT using MTAT v2.1 (https://www.clinicalresearcher.org/software.htm). The threshold cutoff was set at an MEP peak-to-peak amplitude of 50 μV, and the intensity calculated after fifteen pulses was used as the rMT. If no intensity reversals were recorded in the last five pulses, the procedure was restarted.

#### TMS-induced pupil size recordings

For TMS-induced pupil dilation recordings, participants were instructed to maintain fixation within a 5 cm by 5 cm square outlined with a gray (RGB: 128, 128, 128) 0.5 cm border on the centre of a black screen. Live gaze positioning was shown on screen with a gray (RGB: 128, 128, 128) 0.25 cm radius circle. An averaged fixation heatmap for Experiment 2 across all participants is shown in Supplementary Fig. S3. Trials were participant initiated via button press (spacebar using the left hand), with a random jittered baseline duration of either 1.5, 2, or 2.5 seconds, with equal probability (θ*_jitter_*∼0.33). After trial initiation, a TMS pulse was delivered after the jittered baseline duration, followed by a three second delay. As such, trials had a duration of 4.5, 5, or 5.5 seconds. Pupil size was recorded for the whole trial duration. As in Experiment 1, trials were monitored online, and trials contaminated with eye-blinks were replaced, with the exception of eye-blinks accompanying the TMS pulse.

#### Design and procedure

The design and procedure of Experiment 2 were identical to Experiment 1. The only difference from Experiment 1 was that prior to hotspot determination, participants underwent EMG electrode placement and neuronavigation co-registration.

### Experiment 3

The purpose of Experiment 3 was twofold. First, Experiment 3 was designed to investigate whether a CE shift (inhibition) is indeed reflected through TMS-induced pupil dilations, and second, to rule out TMS sensory specific effects, such as pupil responses being due to different skull sensations caused by differences in TMS intensity.

#### Participants

For Experiment 3, 32 participants were initially recruited. However, four participants were excluded from the study. In particular, one participant withdrew due to intolerance of tDCS, one participant was excluded due to reaching the built-in EMG plateau (MEP peak-to-peak ∼ 5 mV) at the baseline condition (pre-tDCS), and two participants were excluded due to inability to collect pupil recordings. As such, 28 participants (12 females, 16 males; age mean = 26.11, *sd* = 6.85, min-max = [18, 49]) were included in Experiment 3. Our sampling plan resembled that of Experiment 2, whereas it included an initial round of our primary analysis (see *Data Analysis* subsection) after collecting data from 20 participants, with a decision threshold of *BF* > 3. Failure to reach the decision threshold resulted in the recruitment of four additional participants before reanalysis. The procedure was repeated, until the decision threshold was reached. Participants that were excluded were replaced. Here, the decision threshold was reached at the third round of analysis (round 1: *n* = 20; round 2: *n* = 24; round 3: *n* = 28).

#### Apparatus

The same set-up used in Experiment 2 was used for Experiment 3. In addition, a battery powered remote mini-CT (Soterix Medical Inc, Woodbridge, USA) transcranial electric stimulation device was used to deliver tDCS. For delivering tDCS, 4.5 cm by 4.5 cm rubber electrodes were placed in 7 cm by 5 cm rectangular sponges, each of which was soaked with 8 ml of 0.9% NaCL saline solution. The procedures for EMG, hotspot determination, rMT estimation, and pupil recordings were identical to Experiment 2. An averaged fixation heatmap for Experiment 3 is provided in Supplementary Fig. S3.

#### Transcranial direct current stimulation

A priming-test tDCS protocol was used to inhibit CE. During tDCS, the anode was placed over the right FDI hotspot (left M1) and the cathode over the contralateral (right) supraorbital region. The electrodes were secured to the scalp using adjustable rubber straps. The protocol comprised 7-minute tDCS with 1 mA current intensity (priming), followed by 5-minute 1 mA tDCS (test), after a 3-minute rest period.

Current intensity was ramp-up and ramp-down controlled over a 10-second duration. When applied alone for at least 10 minutes over M1, the anodal tDCS montage is expected to result in excitatory effects, though, priming anodal tDCS with an initial block of anodal tDCS and a short 3-minute interval in between, reverses the expected anodal stimulation effects, resulting in inhibition [17–19, 33]. The specific protocol was chosen as it has been shown to reliably induce inhibition in healthy participants, even with a single priming-test dose [35], with the inhibitory effects, as reflected through MEPs, peaking approximately 10 minutes post stimulation [19].

#### Design and procedure

Experiment 3 followed a within-participant pre-post design. After EMG set-up, hotspot determination, and rMT estimation, 20 blink free pupil recording trials were collected at an intensity of 120% of rMT (baseline condition). Then the 7-minute priming tDCS block was delivered, followed by a 3-minute rest period. During these 3 minutes, 20 additional blink free pupil recording trials were recorded at 120% of rMT. Recordings from the rest period were only used to control for variability and time dependent effects when pre-processing pupil data (see *Data Pre-Processing* subsection) and were not included in data analysis. The reason that no data analyses were performed for this timepoint is because CE effects following priming are highly variable and often uninformative of the priming-test protocol effects [18, 19, 33]. At the end of the 3-minute rest, the 5-minute test tDCS block was delivered. Twenty blink free pupil recording trials were collected immediately after the test tDCS block (post-0 condition) as well as 10 minutes after the test tDCS block (post-10 condition) at 120% of rMT. This resulted in 80 blink free trials in total (20 trials before tDCS, 20 trials between the priming and test block, 20 trials immediately after the test block, and 20 trials 10 minutes after the test block). An illustration of Experiment 3 is shown in Fig. 2b.

### Data Pre-Processing

#### Pupil data

Pre-processing was implemented for data from the right pupil in Experiment 1, and for both the left and right pupil for Experiment 2 and Experiment 3. Data were monitored online for blinks. Blink contaminated data were marked for exclusion and replaced with new trials. Blinks induced due to TMS (immediately after the TMS pulse) were not excluded. The pre-processing procedure was implemented on the individual level, independently for each participant. In detail, pupil data were first time-locked to the TMS pulse in epochs of 3.5 second duration, ranging from –1 second to 2.5 seconds relative to the TMS pulse (0 seconds). In Experiment 2 and Experiment 3, right and left epoched pupil data were processed both (i) independently and (ii) as an averaged right and left pupil size epoch. After epoching, a window of 75 ms before and 75 ms after (150 ms total) the maximum (Experiment 1) or minimum (Experiment 2 and Experiment 3) value within a 300 ms window post-TMS was interpolated using linear interpolation, to remove TMS induced blink artefacts. After blink interpolation, a *z*-score standardisation was implemented. For each experiment, the *z*-score transformation contained all pupil data for the corresponding epochs (i.e., all right pupil data, or all left pupil data, or all right and left averaged pupil data) from all conditions for each participant, to account for all participant variability and potential time induced change. Absolute *z-*scores above 3 (|*z*| > 3) were transformed to NaN values, to account for possible signal drop. Next, the data were baseline corrected by subtracting the averaged pupil size from the 1 second preceding the TMS pulse across the whole trace, on a trial-by-trial basis. Finally, epochs were separated for each condition, and a mean trace per condition was calculated for each participant, by averaging the 20 trials per condition. For data analysis, peak pupil dilations were calculated as the maximum value within 2 seconds post TMS.

#### Motor evoked potentials

EMG data were recorded in Experiment 2 and Experiment 3, in 300 ms epochs, starting -100 ms prior to the TMS pulse and extending 200 ms after the TMS pulse. Waveforms were baseline corrected by subtracting the averaged activity between - 50 ms to -10 ms relevant to the TMS pulse. For data analysis, we calculated peak-to-peak MEP amplitudes in a window between 17 ms and 45 ms post TMS.

## Data analysis

All data analyses were performed using JASP v0.19.3 (JASP Team, https://jasp-stats.org). Repeated measures ANOVAs were performed using a uniform model prior distribution and Cauchy coefficient priors centered at 0, using a scale *r* = 0.5 for fixed effects and *r* = 1 for random effects. For pairwise comparisons, Wilcoxon signed-rank paired sample *t*-tests were implemented with 1000 samples, using a half-Cauchy prior centered at 0 and a scale of *r* = 0.5. This prior was selected to resemble a minimum difference of interest of approximately *z =* 0.5 [34]. Correlations were performed using a stretched beta prior with parameters α, β = [1, 1], assuming equal probabilities across all possible correlations. Correlation coefficients were calculated using Kendall’s τ-β. Following the results of large simulations [35], a *BF* ≥ 3 (in favor of either model in comparison) was considered adequate evidence to inform our sampling plan analyses.

## Data availability statement

All materials and data used are openly available through the Open Science Framework repository and can be accessed using the following link: https://doi.org/10.17605/OSF.IO/6TYP2.

## Author contributions

PP: conceptualization, methodology, software, formal analysis, investigation, data curation, writing–original draft, visualization, funding acquisition; FM: investigation, writing–review C editing; AM: investigation, methodology, writing–review C editing; NB: methodology, writing–review C editing; AD: investigation, methodology, writing–review C editing; AAA: investigation; XT: investigation; FvE: methodology, writing– review C editing; DAS: conceptualization, methodology, resources, writing–review C editing, supervision; SMS: conceptualization, methodology, resources, writing–review C editing, supervision, funding acquisition.

