## Supplemental Material for "Transcranial magnetic stimulation induced pupil dilations can serve as a cortical excitability measure"

**Supplementary Table S1.** Descriptives and pairwise comparisons of active versus sham TMS condition peak pupil responses in Experiment 1.

| Condition | Mean (*z*-value) | sd | *t*-test $BF_{10}$ |
|---|---|---|---|
| Active 80% | 0.49 | 0.35 | 2.09 |
| Sham 80% | 0.37 | 0.26 | |
| Active 90% | 0.57 | 0.40 | 15.77 |
| Sham 90% | 0.26 | 0.30 | |
| Active 100% | 0.58 | 0.37 | 9.76 |
| Sham 100% | 0.33 | 0.36 | |
| Active 110% | 0.73 | 0.37 | 44.68 |
| Sham 110% | 0.37 | 0.35 | |
| Active 120% | 0.90 | 0.39 | 33.14 |
| Sham 120% | 0.46 | 0.21 | |

**Supplementary Table S2.** Pairwise intensity comparisons of peak pupil dilations per TMS condition in Experiment 1.

| **Measure 1** | **Measure 2** | ***t*-test $BF_{10}$** |
|---|---|---|
| Active 80% | Active 90% | 2.02 |
| Active 80% | Active 100% | 1.29 |
| Active 80% | Active 110% | 21.63 |
| Active 80% | Active 120% | 380.70 |
| Active 90% | Active 100% | 0.52 |
| Active 90% | Active 110% | 12.08 |
| Active 90% | Active 120% | 56.36 |
| Active 100% | Active 110% | 1.88 |
| Active 100% | Active 120% | 115.12 |
| Active 110% | Active 120% | 3.27 |
| Sham 80% | Sham 90% | 0.15 |
| Sham 80% | Sham 100% | 0.26 |
| Sham 80% | Sham 110% | 0.53 |
| Sham 80% | Sham 120% | 1.19 |
| Sham 90% | Sham 100% | 0.80 |
| Sham 90% | Sham 110% | 1.98 |
| Sham 90% | Sham 120% | 46.55 |
| Sham 100% | Sham 110% | 0.54 |
| Sham 100% | Sham 120% | 1.95 |
| Sham 110% | Sham 120% | 1.49 |

**Supplementary Table S3.** Descriptives and pairwise comparisons of active versus sham TMS condition peak pupil responses in Experiment 2.

| Condition | Mean (*z*-value) | sd | *t*-test $BF_{10}$ |
|---|---|---|---|
| Active 80% | 0.36 | 0.30 | 0.61 |
| Sham 80% | 0.32 | 0.20 | |
| Active 90% | 0.44 | 0.44 | 4.46 |
| Sham 90% | 0.30 | 0.24 | |
| Active 100% | 0.50 | 0.40 | 13.22 |
| Sham 100% | 0.30 | 0.23 | |
| Active 110% | 0.56 | 0.42 | 94.01 |
| Sham 110% | 0.29 | 0.18 | |
| Active 120% | 0.65 | 0.42 | 111.45 |
| Sham 120% | 0.38 | 0.31 | |

**Supplementary Table S4.** Pairwise intensity comparisons of peak pupil dilations per TMS condition in Experiment 2.

| **Measure 1** | **Measure 2** | ***t*-test $BF_{10}$** |
|---|---|---|
| Active 80% | Active 90% | 0.83 |
| Active 80% | Active 100% | 17.42 |
| Active 80% | Active 110% | 77.84 |
| Active 80% | Active 120% | 5985.76 |
| Active 90% | Active 100% | 1.16 |
| Active 90% | Active 110% | 18.98 |
| Active 90% | Active 120% | 131.74 |
| Active 100% | Active 110% | 0.34 |
| Active 100% | Active 120% | 4.31 |
| Active 110% | Active 120% | 1.58 |
| Sham 80% | Sham 90% | 0.27 |
| Sham 80% | Sham 100% | 0.23 |
| Sham 80% | Sham 110% | 0.21 |
| Sham 80% | Sham 120% | 0.84 |
| Sham 90% | Sham 100% | 0.37 |
| Sham 90% | Sham 110% | 0.34 |
| Sham 90% | Sham 120% | 2.15 |
| Sham 100% | Sham 110% | 0.25 |
| Sham 100% | Sham 120% | 0.87 |
| Sham 110% | Sham 120% | 1.18 |

**Supplementary Table S5.** Pairwise comparisons of peak pupil dilations between right versus left pupil in Experiment 2.

| **Measure 1** | **Measure 2** | ***t*-test $BF_{10}$** |
|---|---|---|
| Active 80% - Right Pupil | Active 80% - Left Pupil | 0.21 |
| Active 90% - Right Pupil | Active 90% - Left Pupil | 0.25 |
| Active 100% - Right Pupil | Active 100% - Left Pupil | 0.17 |
| Active 110% - Right Pupil | Active 110% - Left Pupil | 0.36 |
| Active 120% - Right Pupil | Active 120% - Left Pupil | 0.14 |

**Supplementary Table S6.** Pairwise comparisons of peak pupil dilations split between Low MEP and High MEP bins following an MEP median split analysis in Experiment 2.

| **Measure 1** | **Measure 2** | ***t*-test $BF_{10}$** |
|---|---|---|
| Active 80% - Low MEP | Active 80% - High MEP | 0.65 |
| Active 90% - Low MEP | Active 90% - High MEP | 0.24 |
| Active 100% - Low MEP | Active 100% - High MEP | 0.23 |
| Active 110% - Low MEP | Active 110% - High MEP | 0.27 |
| Active 120% - Low MEP | Active 120% - High MEP | 1.17 |

**Supplementary Table S7.** MEP and pupil data descriptive statistics from Experiment 3.

| Description | Mean | *sd* |
|---|---|---|
| Baseline MEP peak-to-peak (mV) | 1.52 | 1.29 |
| Priming MEP peak-to-peak (mV) | 1.23 | 1.15 |
| Post-0 MEP peak-to-peak (mV) | 1.41 | 1.38 |
| Post-10 MEP peak-to-peak (mV) | 1.27 | 1.22 |
| Baseline peak pupil dilation (*z*-value) | 0.74 | 0.32 |
| Priming peak pupil dilation (*z*-value) | 0.66 | 0.38 |
| Post-0 peak pupil dilation (*z*-value) | 0.59 | 0.34 |
| Post-10 peak pupil dilation (*z*-value) | 0.63 | 0.32 |

**Supplementary Table S8.** Questions used for study eligibility and safety screening.

| |
|---|
| 1. Have you ever been diagnosed with an eye-condition, such as colorblindness, amblyopia, or strabismus? |
| 2. Do you use corrective lenses (eyeglasses and/or contact lenses)? |
| 3. Have you ever had an adverse reaction to any form of transcranial magnetic stimulation (TMS)? |
| 4. Have you ever had a seizure? |
| 5. Does anyone in your family have epilepsy? |
| 6. Have you ever had an electroencephalogram to measure brain activity (EEG)? |
| 7. Have you ever had a stroke? |
| 8. Have you ever had a serious head injury (including neurosurgery)? |
| 9. Have you ever had any illness that caused brain injury? |
| 10. Do you have any metal in your head (aside from dental work in the mouth), such as shrapnel, surgical clips, or fragments from welding or metalwork? |
| 11. Do you have any implanted devices such as cardiac pacemakers, cochlear implants, medical pumps, or intracardiac lines? |
| 12. Do you wear a hearing aid? |
| 13. Do you suffer from frequent or severe migraines/headaches? |
| 14. Have you ever experienced fainting or blackouts? |
| 15. Have you ever had any other brain-related condition, including any neurological disorder such as Parkinson’s, tics, or stuttering? |
| 16. Do you have any psychiatric conditions such as depression, anxiety, schizophrenia, etc.? |
| 17. Are you taking any medications? |
| 18. Do you suffer from any other medical conditions? |
| 19. Are you pregnant, or is it possible that you may be pregnant? |
| 20. Do you need further explanation of the study’s associated risks? |

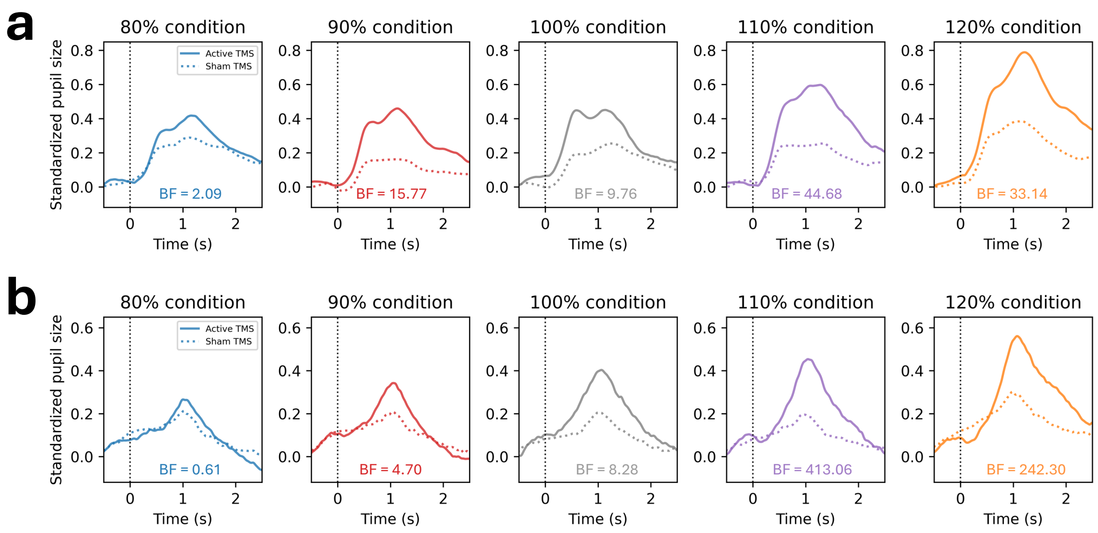


**Supplementary Fig. S1**. Pupil trace comparisons between active and sham TMS in each intensity condition for (**a**) Experiment 1 and (**b**) Experiment 2.

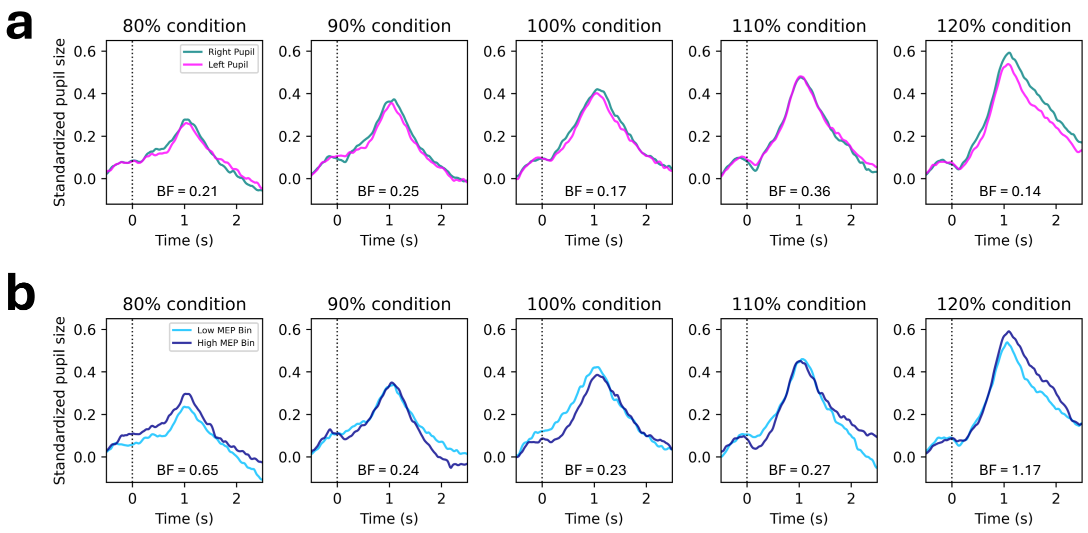


**Supplementary Fig. S2**. Pupil trace comparisons between (**a**) the left and right pupil for each intensity condition and (**b**) between the low versus high MEP trials (identified through median split) for each intensity condition.

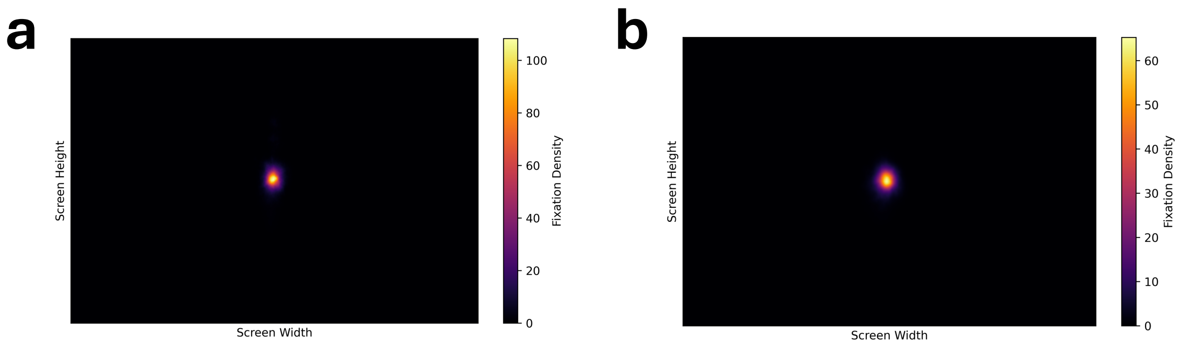


**Supplementary Fig. S3**. Fixation heatmaps averaged across participants for (**a**) Experiment 2 and (**b**) Experiment 3.
